# Integrating detection of copy neutral chromosomal losses in a clinical setting in leukemia and lymphoma by means of allelic imbalance and read depth ratio comparison

**DOI:** 10.1101/590752

**Authors:** Marcus C. Hansen, Oriane Cédile, Maja Ludvigsen, Eigil Kjeldsen, Peter L. Møller, Niels Abildgaard, Peter Hokland, Charlotte G. Nyvold

## Abstract

Chromosomal aberrations are common features of hematological malignancies, with several recurrent aberrations recognized as important diagnostic or prognostic molecular markers. While genome-wide genotyping and genomic hybridization microarray analyses have been implemented in clinical laboratories for several years the usage of next generation sequencing for detection of acquired copy number alterations has not yet reached full clinical integration. One evident problem is the identification of copy neutral loss of heterozygosity (CN-LoH), which is not detectable by sequencing read depth correlation or the analogous microarray CGH. We selected 23 paired samples of hematological disorders from 14 individuals, focusing on leukemia and lymphomas, and tested whether a low complexity approach, relying on Fisher’s exact test or χ^2^, is efficient for the analysis of variant allele frequencies with read-depth ratio correlations in order to resolve both copy-altering and neutral chromosomal aberrations. This combination helped to identify 69 altered chromosomes and offered mutual confirmation. Moreover, six CN-LoHs (>1% AF shifts, p<0.01) were found with one additional suspected low frequency deletion (~10-15% burden) in a case of CLL. We conclude that this simple method is directly clinically applicable for the detection of copy neutral chromosomal loss of single genes from WES with intermediate coverage.

**Availability:** Mentioned plot software is available at Harvard Dataverse http://doi.org/10.7910/DVN/KFMGNY

**Supplementary information:** Supplementary plots are available

## 1 Introduction

Chromosomal instability and cellular aneuploidy is a common feature of solid tumors and hematological malignancies. Several recurrent aberrations involving copy changes are recognized as important diagnostic or prognostic features, such as with monosomy 7 or 7q deletions in myeloid neoplasms and 3q or 8q gains in subtypes of lymphoid neoplasms. While genome-wide genotyping and genomic hybridization microarray analyses have been implemented as diagnostic tools in clinical laboratories, the usage of next generation sequencing (NGS) for detection structural variants, such as acquired copy number alteration (CNA), has not yet reached full maturation and its robustness and algorithms for routine clinical application is still being evaluated.

Several algorithmic strategies based on whole exome sequencing (WES) have been proposed, primarily focusing on read depth alterations affected by the chromosomal imbalance of CNA. Some studies have, however, demonstrated a discouraging degree of concordance between developed approaches (Hong et al. 2016; Zare et al. 2017), while copy neutral events escape detection by the majority of tools. One study concluded that the small overlap is symptomatic of an immature WES-based CNA detection (Yao et al. 2017). Among other problems, low coverage is of concern and affects both read depth ratios and the dispersion of variant allele frequencies. It is often unclear how the different algorithms perform under such situations. A seminal study showed that the number of false positives are highly dependent on the size of the aberrations, where only a few percent of CNAs spanning one or two exons can be confirmed by array CGH (Retterer et al. 2015). This figure rises to 89% for deletions and 60% for duplications, when spanning three or more exons. Retterer and colleagues suggested that WES, which overall has not changed notably since the early years, was not mature to replace arrayCGH and SNP arrays (Retterer et al. 2015). Batch effect was another suspected cause of variation or incompatibility in this paper. We now know this to be an inevitable problem, when correlating samples from different sequencing runs. Taking these matters into account, the detection of CNA calls for demonstration of algorithms for clinical implementation with a high degree of methodological transparency – offering at least a satisfactory minimum viable product for the clinic. In addition, a present issue in contemporary cancer cytogenetics is the practical identification of copy neutral loss of heterozygosity (CN-LoH) or isodisomy, which is the aim of this proof-of-concept paper.

Both the CNAs and CN-LoHs are types of structural variants, however the precise definition of such aberrations is not entirely fixed, let alone the usage of the term copy number variation. Regarding size, a threshold of one kilobase has arbitrarily been suggested (Ionita-Laza et al. 2008; Redon et al. 2006), which may reflect the methodological resolution (Zhang et al. 2009) rather than representing biological meaning. On a macromolecular scale the copy number change can perhaps more objectively be defined as the discrete change of the single gene, with a typical coding length in the order of a few kilobases (H. sapiens Annotation Release 108, NCBI, MD, USA), leading to and increase or decrease in protein products or activity hereof by means of transcriptional abundancy, haploinsufíïciency, phenotypic penetrance etc.

Copy neutral losses can arise from partial or complete loss of a homologous chromosome, but also occur as a secondary effect from trisomy or monosomy. In these cases, the disomic state may be ‘rescued’, and thus not lead to cellular aneuploidy in its traditional sense. This seemingly common event (O’Keefe et al. 2010), along with its potentially immediate clinical consequences, cannot be appreciated by means of arrayCGH alone, nor is it retrievable by sequencing read depth correlation. Instead, read depths are complemented by variant allele frequencies (AF), analogous to that of single nucleotide polymorphism microarrays and comparative genomic hybridization in combination (Wiszniewska et al. 2014). Likewise, it is not possible to resolve a stretch with loss of heterozygosity as a frank deletion or a copy neutral event from AF alone. Hence, robust assays and algorithms represent crucial steps in moving cytogenetics towards NGS cytogenomics. Meanwhile, with a decade passed since the introduction of exome sequencing (Choi et al. 2009; Ng et al. 2009), arrays are still often the used methodology in contemporary studies, as exemplified by Rego et al. (Rego de Paula Junior et al. 2018).

In this paper we address two subsequent problems: **1)** general detection of CNAs by exome sequencing by transparent and simple statistics and signal processing in order to perform **2)** efficient differentiation of copy neutral events in a diverse set of hematological malignancies, not previously investigated for copy neutral deletions by conventional cytogenetics. We hypothesized that Fisher’s exact and χ^2^ is sufficient for detection of these structural variants by means of control-paired variant allele frequencies in medium to high quality whole exome sequencing. Not only do juxtaposed gene-wise read depth ratios confirm the majority of these frequency changes as arising from acquired CNA, but in these cases also rule out the presence of a CN-LoH. The clear motivation is to establish a simple methodological approach for the detection of copy neutral deletions in hematological malignancies, which is independent of neoplastic subtype and input material and clinically applicable. In addition, we elaborate on the fundamentals and assumptions of AF and the highly relevant concept of molecular burden in the latter part of the paper. Here, we position CN-LoH as a special case of CNAs, as it arises as a variation from the normal copy state of the autosomes with one paternal and one maternal copy. Consequently, we use the term to cover both structural variants in the following for practical reasons, if not explicitly stated otherwise.

## 2 Methods

Sequencing analysis was based on WES from 14 individuals, selected to represent different hematological disorders with no clinically assumed CNAs (acute myeloid leukemia with normal karyotype (CN-AML, 5), monoclonal B-cell lymphocytosis (MBL, 1), intermediate ((AML, 1), chronic lymphoblastic leukemia (CLL, 1), T-cell acute lymphoblastic leukemia/ETP-ALL (T-ALL, 1), chronic myeloid leukemia (CML, 1)) to frequent complex copy alterations (mantle cell lymphoma (MCL, 4)). In total, 38 samples had been exome sequenced (Illumina HiSeq 2000/2500), constituting 23 control-paired sample sets, of which 14 samples were diagnostic samples in addition to 9 relapse samples. Control samples (15) consisted of remission, skin samples or non-malignant B- or T-cells (see supplement). Material for library preparation was derived from sorted cell populations or unsorted cells (bulk). Library preparation was performed with SureSelect (Agilent Technologies, CA, USA) or Nextera (Illumina, CA, USA) kits.

Alignment (GRCh37) and variant calling was performed using BWA (Li and Durbin 2009) and GATK (McKenna et al. 2010) (GATK, version 3.6/3.8) workflow (Van der Auwera et al. 2013). Variants and reads located outside targeted regions were excluded from the analyses, and a base quality threshold of 25 and minimum allele read depth 20 was applied. Gene-wise read depths were retrieved with bedtools (Quinlan and Hall 2010) from RefSeq coordinates (UCSC, (Karolchik et al. 2004)). Fisher’s exact test was used to detect allelic imbalance. The significance level was set as 0.01 based on the total number of chromosomes investigated and the threshold of single nucleotide variant (SNV) allele frequency shifts in order to define it as a structural variant (>1% per chromosome) – weighted against tolerable number of false positive calls. Test of change in reference and alternate allele read depths in sample versus control sample was complemented by the Brown-Forsythe test for equal variances (Brown and Forsythe 1974) to detect low burden copy changes on chromosomal level. Statistics and data analysis of variants (in variant call format) and gene read depth (BED format) were performed in Mathematica (Wolfram Research, Ill, USA). Gene-wise DP ratios were compared to the established CNA tool VarScan2 (Koboldt & al, 2012), which evaluates read depth ratios. Visual representations of the results were prepared for publication in Prism 7 (GraphPad Software, CA, USA) and Mathematica. We developed a general-purpose plotting software for the juxtaposition of AFs and DP ratios, implementing both Fisher’s exact and χ^2^ (*see Availability section*).

## 3 Results

### Quality control of sequenced samples

Sequencing yield spanned 48.5–188.6 million (median 84.3×10^6^) paired-end reads mapped to the human genome (mapping efficiency > 99%). The resulting 23 paired variant call sets (38 samples in total) comprised a median of 19,101 coding variants, matching the expected number (Frebourg 2014; Ng et al. 2009), with a median (Q2) read depth of 96 and lower (Q1) and upper quartiles (Q3) at 71 and 136, respectively. Whereas the mean chromosomal heterozygous allele frequencies (0.2 < AF < 0.8) was found to be constant (Fig. 1, Q[1,2,3] = [0.478, 0.482, 0.486]), with few outlier chromosomes (n=7), the chromosomal AF standard deviations (SD) were generally comparable (Q[1,2,3] = [0.062, 0.068, 0.074]), while comprising a large number of outliers (49, Fig. 1) arising from allelic imbalance. 96% of the autosomes was found Gaussian with regards to AF distributions of heterozygous variants (805/836 at 99.9% confidence level, Anderson-Darling test).

**Figure 1).**
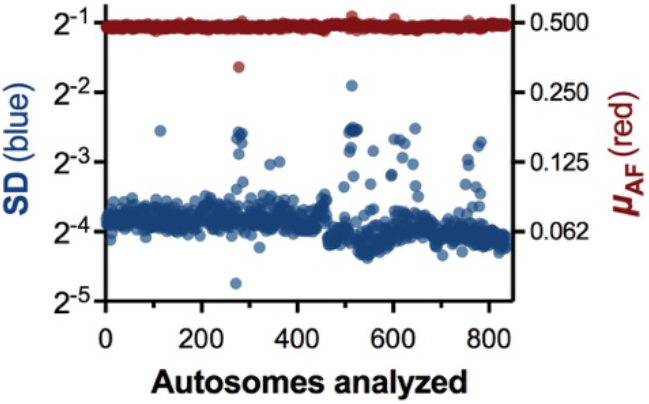
Fluctuations of the standard deviation (SD). Whereas the SD (blue) of heterozygous allele frequencies (0.2 < VAF < 0.8) was comparable between chromosomes unaffected by CNVs of same sequencing batch, the introduction of a copy gain or low frequency copy loss increased the SD. The mean chromosomal heterozygous allele frequency (red, μ_AF_) did not differ to any significant extent. The AF symmetry of, e.g., a trisomy left the mean unaltered,

Gene-wise read depth (DP) pairing of target and control samples revealed high correlation coefficients (Spearman *ρ*) with a median of 0.99. Whereas samples devoid of CNAs displayed a single linear appearance (Fig. 2A), compared to samples with copy changes (Fig. 2B). Correlation coefficients were sensitive towards dissimilar read depths, while CNAs did not affect *ρ* to same extend due to symmetry, and thus represented a reasonable quality measure. Two samples from the same patient (CML) were excluded due to batch effects.

**Figure 2).**
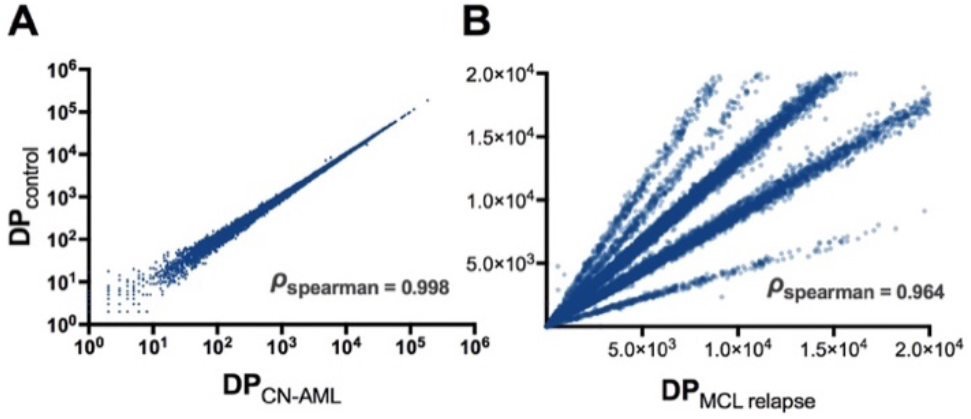
Correlation of read depths from paired samples. Gene-wise correlation between cytogenetic normal sample and paired control shows a single linear relationship for samples run in same sequencing batch (A, log-scaled). In contrast, tumor samples with copy alterations show discrete shifts, which reflect the copy state and size, here exemplified by mantle cell lymphoma relapse with 5 discrete copy states (B).

### Read depth paired allele frequency changes

In the calculation Fisher’s exact test was implemented to detect CNAs by allelic imbalance in exome sequenced hematological neoplasms versus disease free samples. The selected significance level (α=0.01) and a threshold of more than one percent significant allele frequency shifts led to the initial output of 68 significant chromosomes with suspected allelic imbalance with several chromosomes affected by both copy gain and loss. Four chromosomes were visually recognized as false-positives, resulting from random AF shifts in regions otherwise devoid of copy change. In contrast, five chromosomes did not pass the test because of a low number of significantly affected variant alleles. These was readily resolved from cross-check of DP ratios. Thus, the systematic comparison of altered allele frequencies with corresponding read depth change lead to detection of 69 affected chromosomes in total, of which 6 significant CN-LoH (p < 0.01, n > 1%) were detected (See supplement). In addition, a low frequency deletion of chromosome 13 (Table 1*, CLL, app. 10–15%), not passing the selected threshold of 1% altered variants and alpha level (14/230 significant variants, p_mean_ < 0.02), was suspected. Robust test of equal variances (Brown and Forsythe 1974) between the paired diagnosis and remission sample revealed an extreme p-value of 1.7×10^−11^ in the latter sample in contrast to the other autosomes (p_median_ = 0.019) with possible applicability to complement Fisher’s exact test in cases with low burden. This approach was motivated by the distinct fluctuation of standard deviations of affected chromosomes (Fig. 1). The resulting findings are summarized in Table 1 and the supplement. The other CN-LoHs included high-burden (>90%) acquired isodisomies (Fig. 3A), partial deletions (3B) and a lower burden isodisomy in a case of CN-AML (approximately 20%, 3C).

**Table 1).**
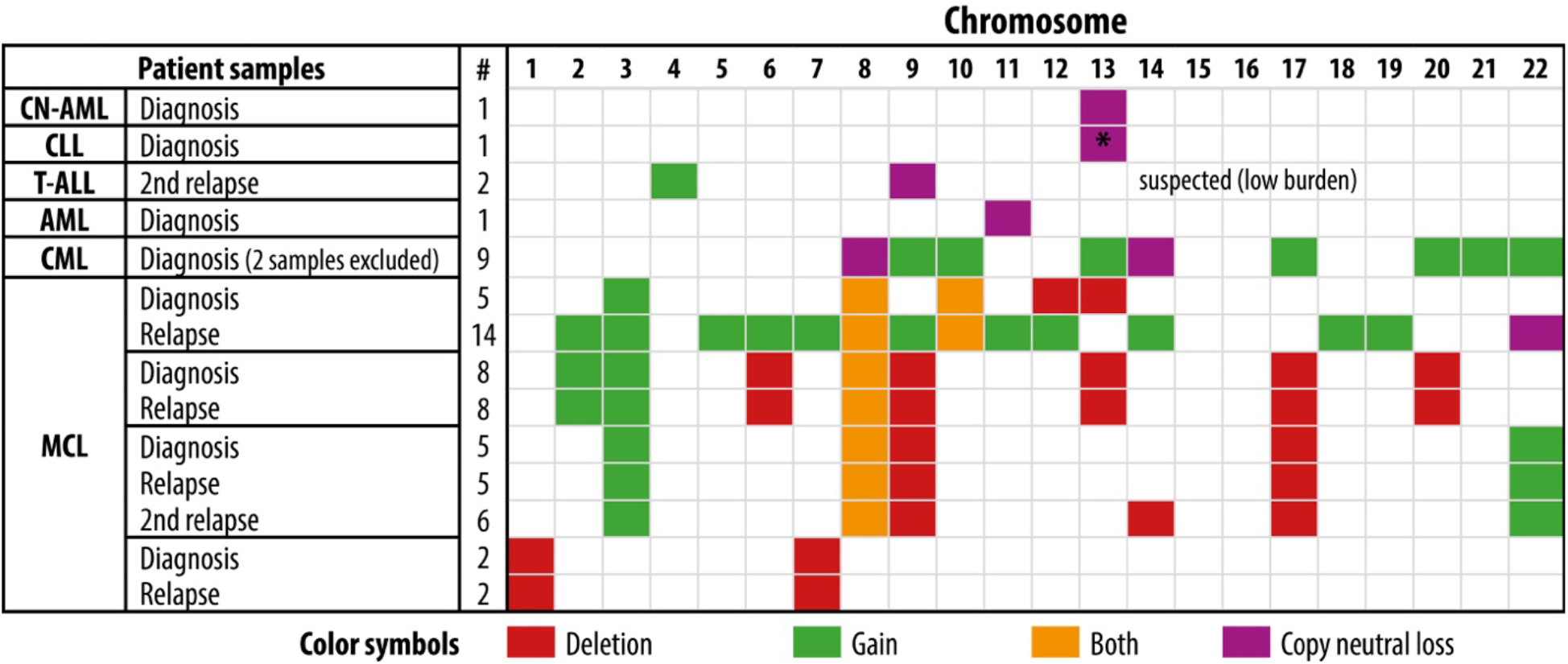
Overview of CNV and CN-LoH.

**Figure 3).**
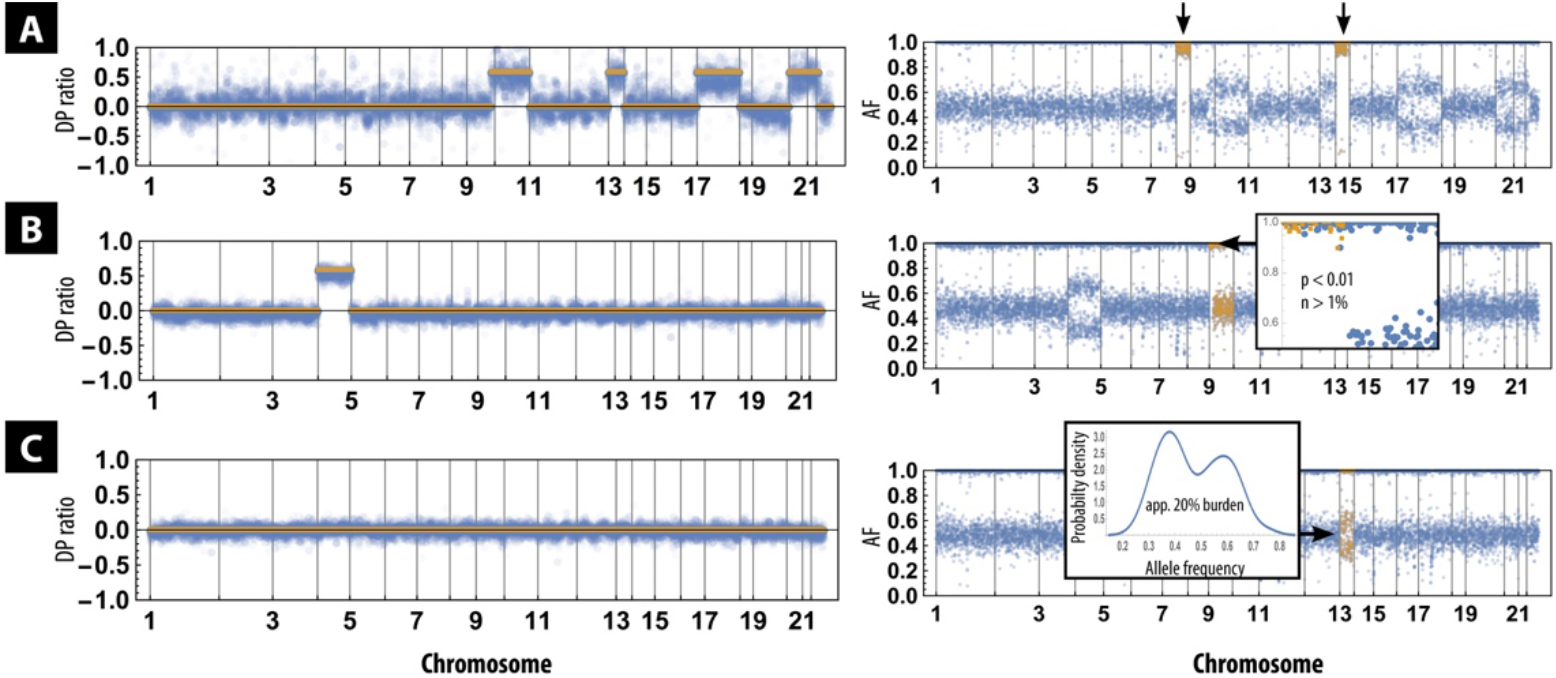
Three examples of allele frequency (AF) paired read depth (DP) ratios. The CML sample shows two acquired isodisomies on chr 8 and 14 (A, right arrows). While the DP ratio is heavily influenced by noise from sequencing batch effect (left), the AF itself is not affected (right). A partial chromosomal loss is shown in B (T-ALL), with a high resolution in gene-wise DP ratio, also confirming trisomy 4. Lower burden CN-LoH (CN-AML, C) represents a problem within detection of structural variants may fail to reach statistical significance in any algorithm.

## 4 Discussions

Detection of increase or decrease in the number of chromosomes is an important part of cytogenetics in hematological malignancies and other types of cancer. The motivation behind this paper was to extend the general detection of CNAs by exome sequencing to the subtle distinction of copy-neutral chromosomal losses by practical means for direct clinical integration. The relevance is justified by the fact that current sequencing read depth correlations, of which many tools exist, do not detect this type of deletions. Some combined tools are available, but the applicability generally lacks transparency. From this proof-of-concept study, we correlated CNA detection from exome sequences by two different, reproducible methods: detection of allelic imbalance by AF and the read depth ratio of a given gene in order to directly pinpoint CN-LoHs.

All escaping previous clinical cytogenetic analyses, the cancerous samples from 6 of the 14 patients were found to harbor 7 different copy neutral deletions in total. One of these was detected in a diagnostic sample from a patient otherwise diagnosed as CN-AML. As expected, evaluation of read depths alone did not reveal these specific aberrations, emphasizing the necessity of implementing AF analysis. We found that smoothing of read depth ratios, such as with a median filter, is indispensable when assessing samples of different signal-to-noise ratios.

Now, as NGS is being utilized widely in laboratory diagnostics, it is of utmost importance to gain an accurate detection of CN-LoH adding substantial value to current cytogenetics. From our results, we suggest that pre-sequencing, such as correct sampling, cell sorting and sequencing parameters, most notably coverage and batch effects, are as critical steps as the different algorithms available. We also observed that heterozygous AF standard deviations are influenced by the total number of batch reads and that CNA has a profound effect on the AF distribution itself: Large deviations in AF, also utilized by SNP arrays, are often visible by frequency plots alone. This allelic imbalance is readily comprehensible as the observed distribution is directly influenced by copy gain and loss. Assessment of coverage enables more sensitive detection and parallel analyses of both offer stringent validation of CNAs and resolution of partial CN-LoH or complete isodisomies. A very close correlation between CNAs detected by means of AF by Fisher’s exact test and change in DP ratio exists, as shown here. Also, we observed that statistical comparison of heterozygous AF standard deviations is a potentially powerful supplement for the detection of low-burden chromosomal allelic imbalance.

While specific copy number changes can be *suspected* from sample allele frequencies *without a paired control*, this is not feasible by coverage analysis. Additional caveats include batch variability, which affects AFs to a lesser degree. Thus, derivation of chromosomal copy alterations from variant AF is a powerful tool on its own. Although the most frequent utilized methods rely on DP ratios the usage of AF has been described several years ago (Wang et al. 2015).

Common for the approaches presented is the theoretical ability to evaluate the tumor burden of the copy change, which will not be elaborated extensively here. One important notice is that the distributions from heterozygous alleles affected by CNA may suffer from allele frequency overlap as a function of both variance and burden. Theoretically, the frequency maxima of the two distributions must be separated by more than 2σ at lower burden to resolve a bimodal allele frequency shift (Schilling et al. 2002). Conceptually, the direct use of allele frequencies in a clinical laboratory context can be appreciated by the following deduction, where the allele frequency signal reflects a mosaicism of diploid normal cells a malignant clone with a discrete copy number state: In cytogenically normal samples the mean chromosomal allele frequency of heterozygous variants approximates one-half (Fig. 4, Eq. 1). From this simple model the general extension describing altered copy state can be expressed as Eq. 2 (Fig. 4).

**Figure 4).**
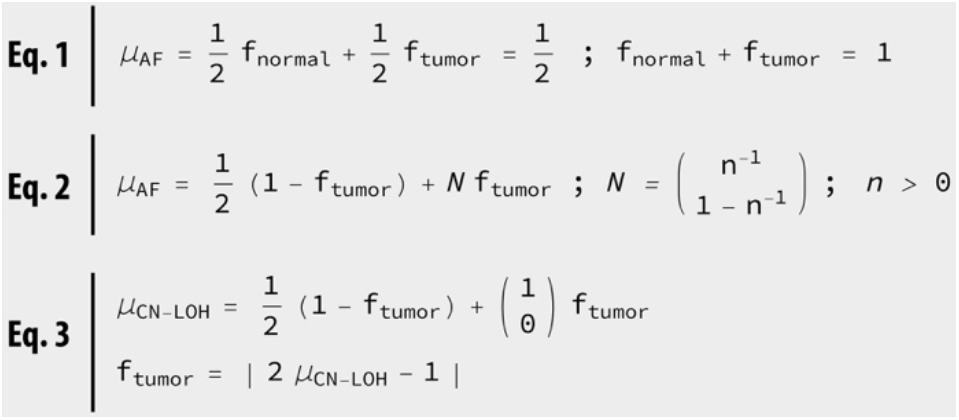
Calculation of mean allele frequencies and burden. A mosaic with a fraction normal (f_normal_) and tumor cells (f_tumor_) with normal diploid karyotype will show a mean variant allele frequency of one-half (Eq. 1). This simple relationship can be generalized to aneusomy of n copies (Eq. 2). Hence, in the case of a copy neutral event (CN-LoH) the burden can be derived and estimated (Eq. 3).

It follows that the mean allele frequencies, *μ*AF, will be symmetrically distributed around one-half as a consequence of reference and alternate alleles in the variant sets. Once a specific chromosomal copy state has been suspected, such a monosomy or isodisomy, the burden can be estimated for this state (Fig. 4, Eq. 3). As pointed out, resolution is poor in samples with a low burden and calls for more advanced mathematical derivations not covered here. These methods have obvious limitations: exome sequencing is inherently noisy, with several contributions to the signal-to-noise ratio – such as coverage and target capture bias. It is not possible to firmly determine copy number from AFs alone, and it can easily be derived from above that a monosomy of one-third burden presents itself equal to a full burden trisomy in terms of allelic imbalance. This is not practically a problem, when dealing with CN-LoH here. Multiple clones can, and often do, exist at time of sampling. Not only do subclonal CNAs make detection more difficult, but also affect the estimation of cancer burden. In addition, it is generally not possible to predict whether the copy loss or gain is a result of an unbalanced translocation, nor is it feasible to detect balanced translocations with exome sequencing using short reads.

Development of practical methods for identifying structural variants is an ongoing area of research, with some indications that obtaining stable results from exome sequencing requires strict guidelines. As simple algorithms can be adequate for detecting CNA we evaluate that sample handling and a proper laboratory setup is equally important for successful detection. Currently, NGS and bioinformatics may not be ready to replace other cytogenetics laboratory modalities. Nonetheless, it is a source of robust methods to fortify or complement existing analyses, such as in the matter of detecting copy neutral chromosomal variation. In order for NGS to succeed in clinical decision making transparent tools and techniques are needed in the transitional phase between research and applied routine detection. Many recent papers dealing with structural variants are devoid of usage of variant AF, perhaps because of this lack of transparency and the general discrepancy in CNA detection. An otherwise excellent review by (Hehir-Kwa et al. 2015), published when exome sequencing and CNA detection gained momentum, lists four different methods of CNA detection, while the potential usage discussion of AF is left untouched – in spite of the development of SNP arrays the previous years.

A detailed estimation of the frequency of acquired CN-LoH in hematological malignancies remains to be revealed. However, it can be concluded from the results and the existing literature that these aberrations may be frequent, even in CN-AML, and thus cannot be neglected. We also conclude that simple statistics on the basis of allele frequency shifts, juxtaposed with DP ratios, is robust for the detection of copy neutral losses if sampling and sequencing quality meets certain criteria. This has tremendous value, but pose several problems as described. If control samples are used for the purpose of DP ratio analyses, these must be sequenced simultaneously to avoid sequencing batch effects.

## Acknowledgements

MCH wish to thank research colleagues at AUH and OUH and MD Mads O.B. Lorenzen for discussions.

## Availability

The methodology has been ported to a standalone plotting application under OS X freely available at Harvard Dataverse http://doi.org/10.7910/DVN/KFMGNY for demonstration purposes, where gene-wise read depth ratios and variant allele frequencies are juxtaposed. Simple χ^2^ (optionally Fisher’s exact test) statistics are performed on control-paired allele frequencies.

