## Supplemental data for "Integrating detection of copy neutral chromosomal losses in a clinical setting in leukemia and lymphoma by means of allelic imbalance and read depth ratio comparison"

*Table S1) Expanded overview of CNV and CN-LoH*

| Patient samples | | | | # | Chromosome 1 | 2 | 3 | 4 | 5 | 6 | 7 | 8 | 9 | 10 | 11 | 12 | 13 | 14 | 15 | 16 | 17 | 18 | 19 | 20 | 21 | 22 |
|---|---|---|---|---|---|---|---|---|---|---|---|---|---|---|---|---|---|---|---|---|---|---|---|---|---|---|
| **Cytogenetically normal acute myeloid leukemia, pt. 1** | Diagnosis | BM | Remission | | | | | | | | | | | | | | | | | | | | | | | |
| **Cytogenetically normal acute myeloid leukemia, pt. 2** | Diagnosis | BM | Remission | 1 | | | | | | | | | | | | | Copy neutral loss | | | | | | | | | |
| **Cytogenetically normal acute myeloid leukemia, pt. 3** | Diagnosis | BM | Remission | | | | | | | | | | | | | | | | | | | | | | | |
| **Cytogenetically normal acute myeloid leukemia, pt. 4** | Diagnosis | BM | Remission | | | | | | | | | | | | | | | | | | | | | | | |
| **Cytogenetically normal acute myeloid leukemia, pt. 5** | Diagnosis | BM | Remission | | | | | | | | | | | | | | | | | | | | | | | |
| **Chronic lymphocytic leukemia** | Diagnosis | PB | Sorted T-cells | 1 | | | | | | | | | | | | | Copy neutral loss * suspected (low burden) | | | | | | | | | |
| **Monoclonal B-cell lymphocytosis** | Surveillance | PB | Sorted T-cells | 0 | | | | | | | | | | | | | | | | | | | | | | |
| **T-cell Acute Lymphoblastic Leukaemia** | Diagnosis | PB | Fibroblasts and keratinocytes (2 samples) | 0 | | | | | | | | | | | | | | | | | | | | | | |
| | 1st relapse | PB | | 0 | | | | | | | | | | | | | | | | | | | | | | |
| | 2nd relapse | PB | | 2 | | | | Gain | | | | | Copy neutral loss | | | | | | | | | | | | | |
| **Acute myeloid leukemia** | Diagnosis | BM | Remission | 1 | | | | | | | | | | | Copy neutral loss | | | | | | | | | | | |
| **Chronic myeloid leukemia** | Diagnosis (excluded) | PB | Remission | - | | | | | | | | | | | | | | | | | | | | | | |
| | Blast crisis | PB | | 9 | | | | | | | | Copy neutral loss | Gain | Gain | | | Gain | Copy neutral loss | | | Gain | | | Gain | Gain | Gain |
| | CNS relapse (excluded) | CSF | | - | | | | | | | | | | | | | | | | | | | | | | |
| **Mantle cell lymphoma, pt. 1** | Diagnosis | Sorted B-cells | Sorted T-cells | 5 | | | Gain | | | | | Both | | Both | | Deletion | Deletion | | | | | | | | | |
| | Relapse | | | 14 | | Gain | Gain | | Gain | Gain | Gain | Both | Gain | Both | Gain | Gain | | Gain | | | | Gain | Gain | | | Copy neutral loss |
| **Mantle cell lymphoma, pt. 2** | Diagnosis | Sorted B-cells | Sorted T-cells | 8 | | Gain | Gain | | | Deletion | | Both | Deletion | | | | Deletion | | | | Deletion | | | Deletion | | |
| | Relapse | | | 8 | | Gain | Gain | | | Deletion | | Both | Deletion | | | | Deletion | | | | Deletion | | | Deletion | | |
| **Mantle cell lymphoma, pt. 3** | Diagnosis | Sorted B-cells | Sorted T-cells | 5 | | | Gain | | | | | Both | Deletion | | | | | | | | Deletion | | | | | Gain |
| | Relapse | | | 5 | | | Gain | | | | | Both | Deletion | | | | | | | | Deletion | | | | | Gain |
| | 2nd relapse | | | 6 | | | Gain | | | | | Both | Deletion | | | | | Deletion | | | Deletion | | | | | Gain |
| **Mantle cell lymphoma, pt. 4** | Diagnosis | Sorted B-cells | Sorted T-cells | 2 | Deletion | | | | | | Deletion | | | | | | | | | | | | | | | |
| | Relapse | | | 2 | Deletion | | | | | | Deletion | | | | | | | | | | | | | | | |

**Color symbols** Deletion Gain Both Copy neutral loss

### Cytogenetically normal acute myeloid leukemia

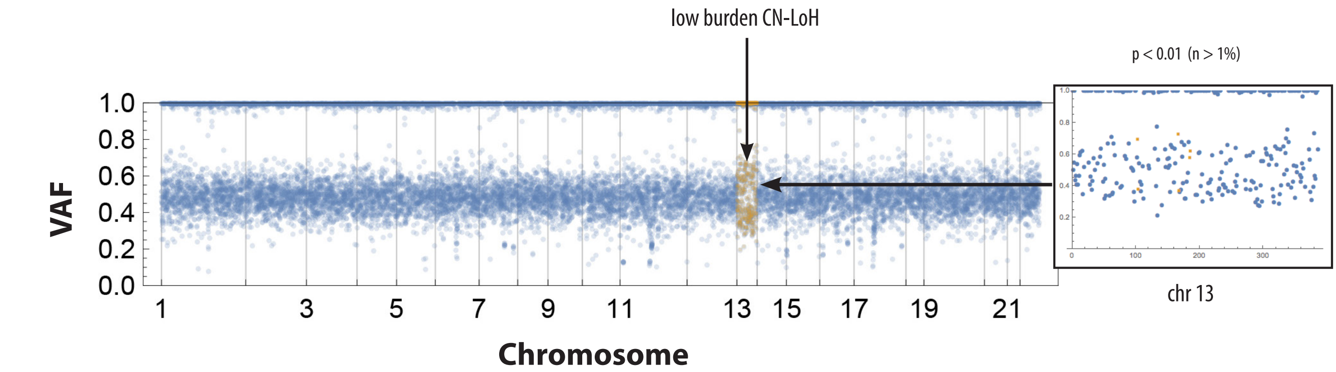


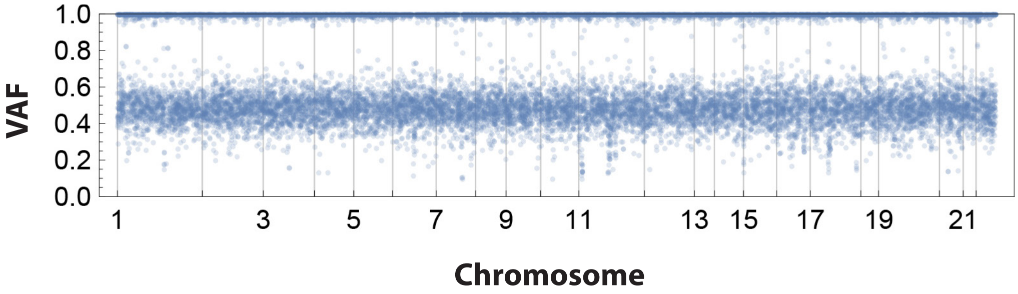


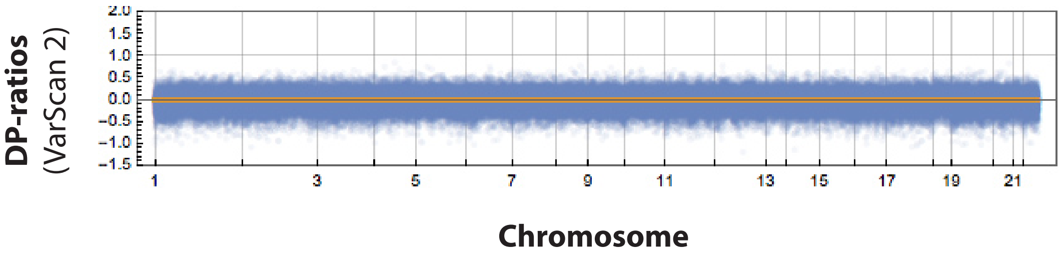


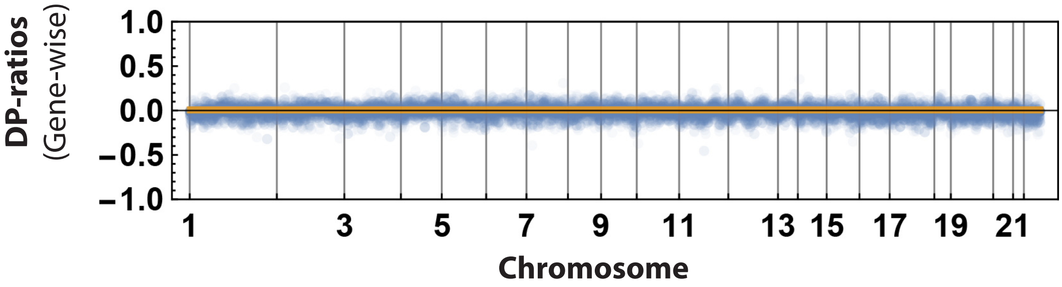

### Chronic lymphocytic leukemia

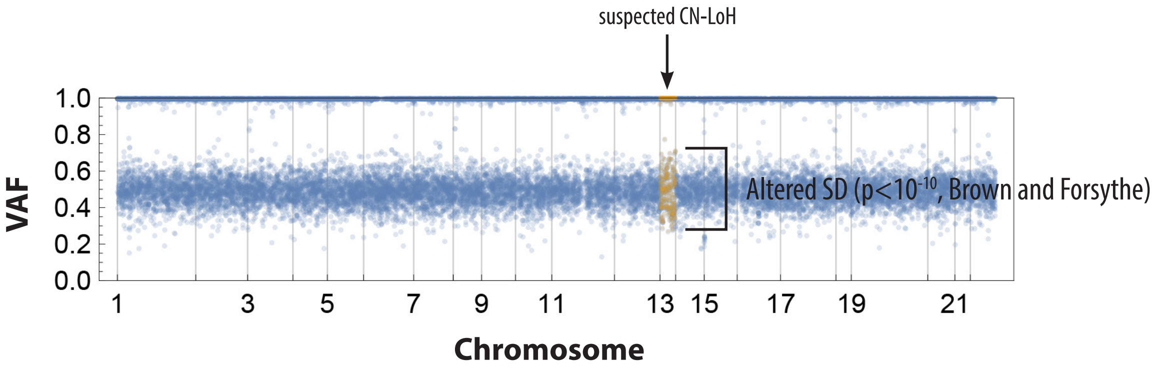


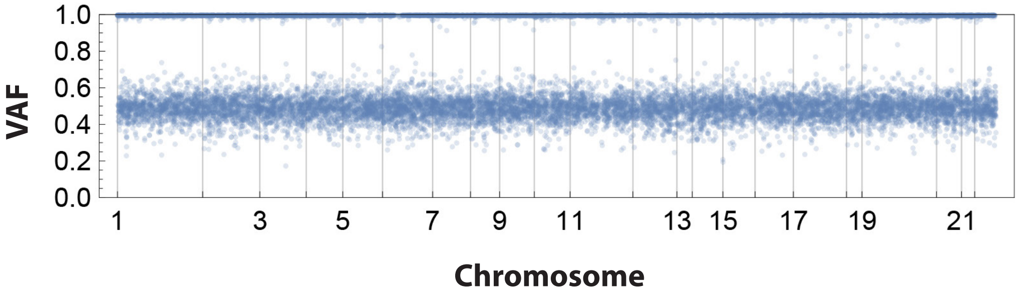


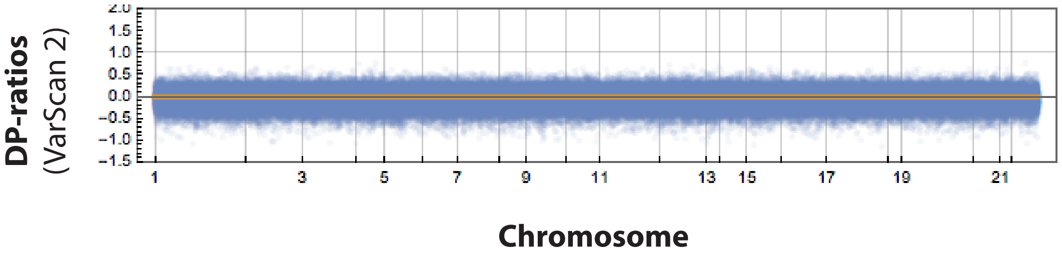


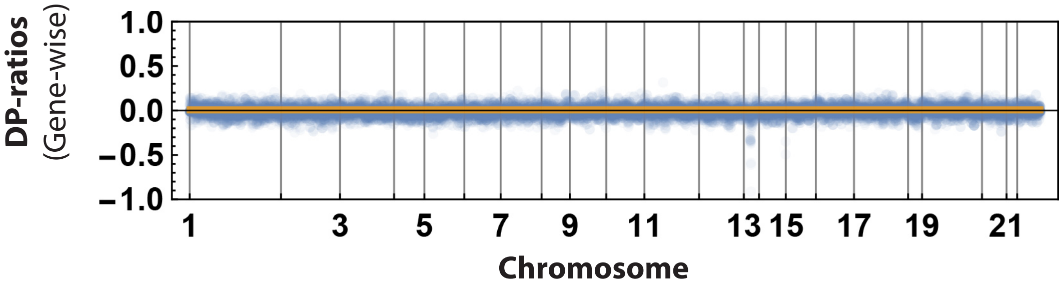

### Mantle cell lymphoma

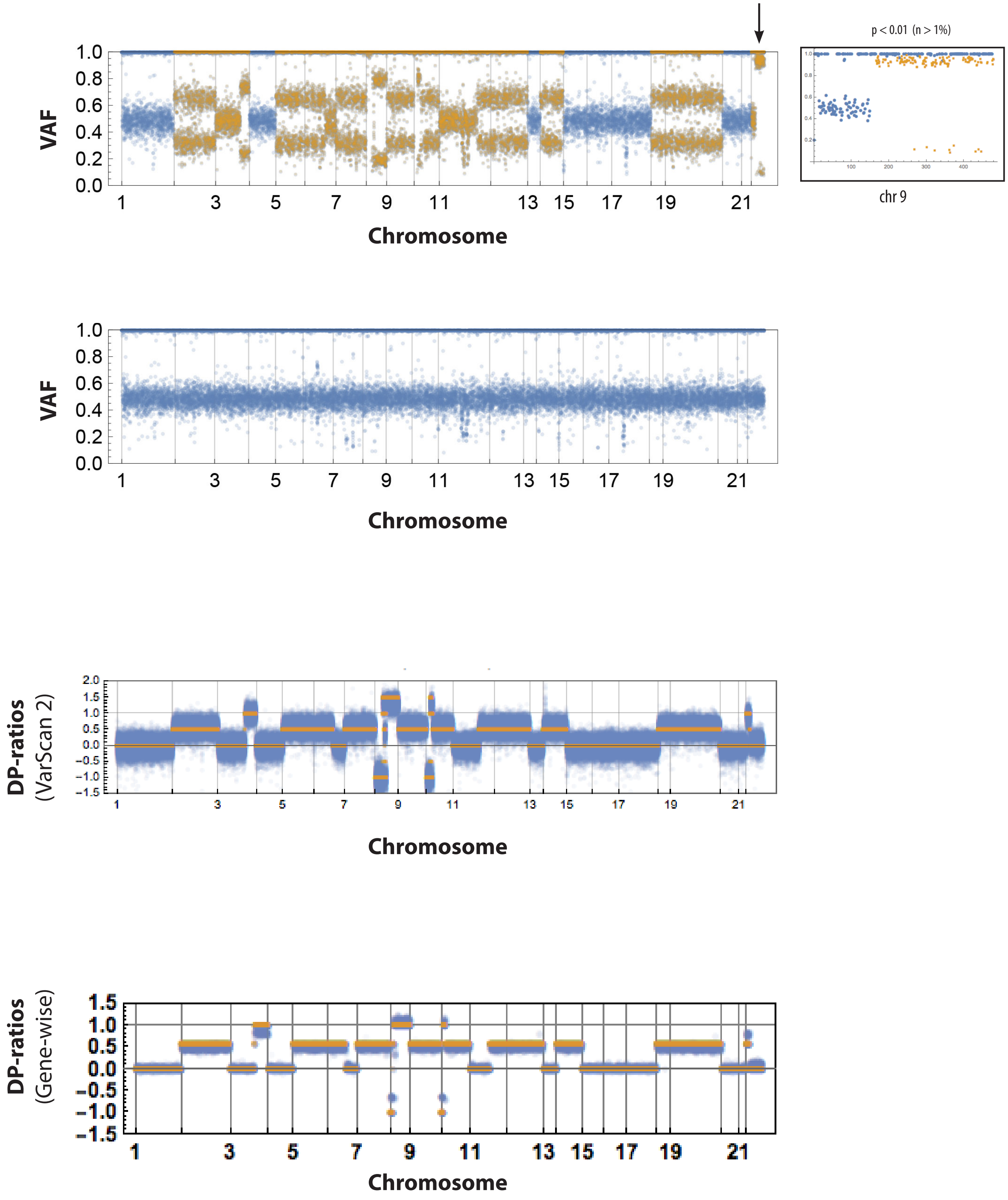

### T-cell acute lymphoblastic leukemia

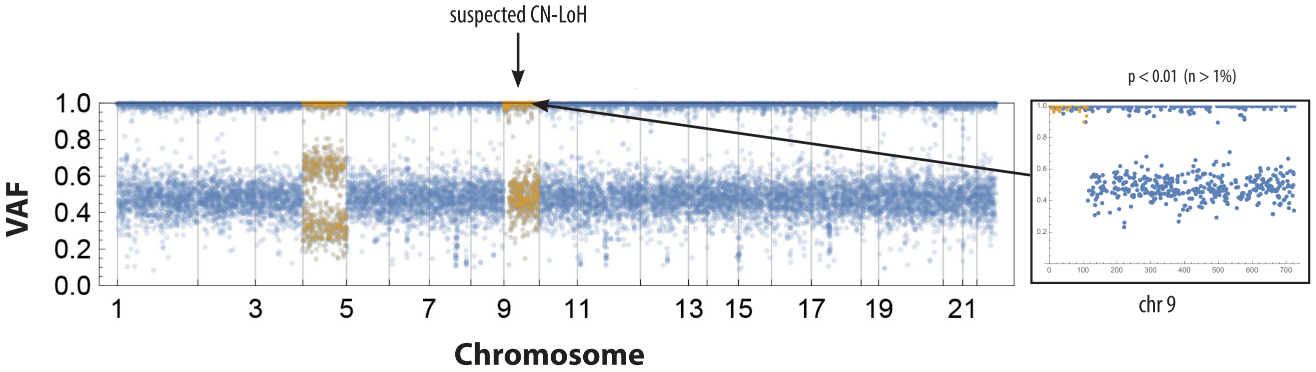


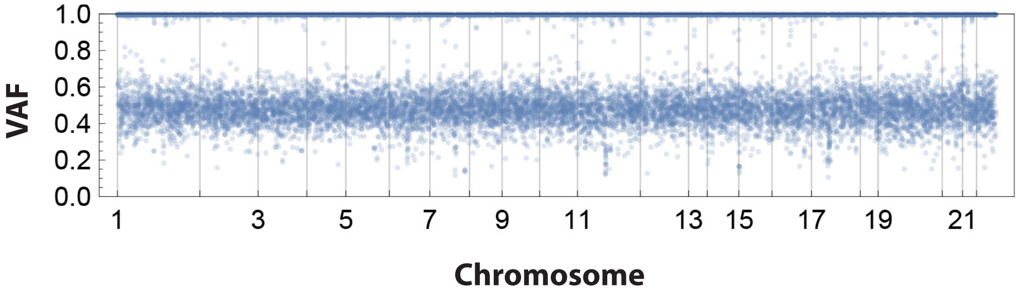


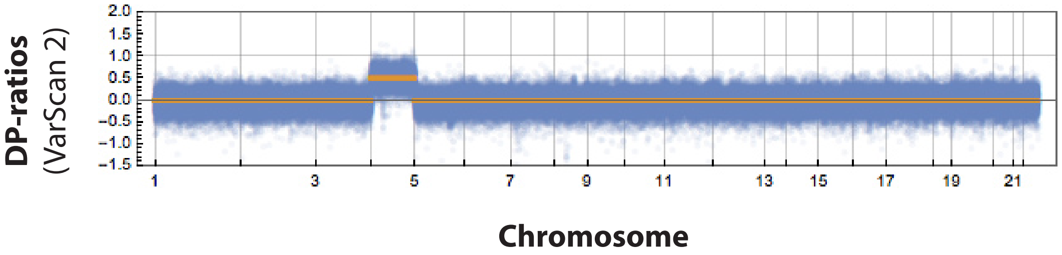


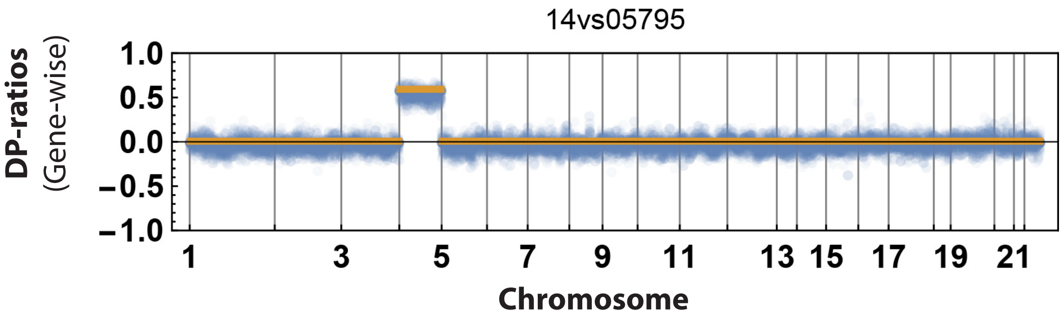

### Chronic myeloid leukemia blast crisis

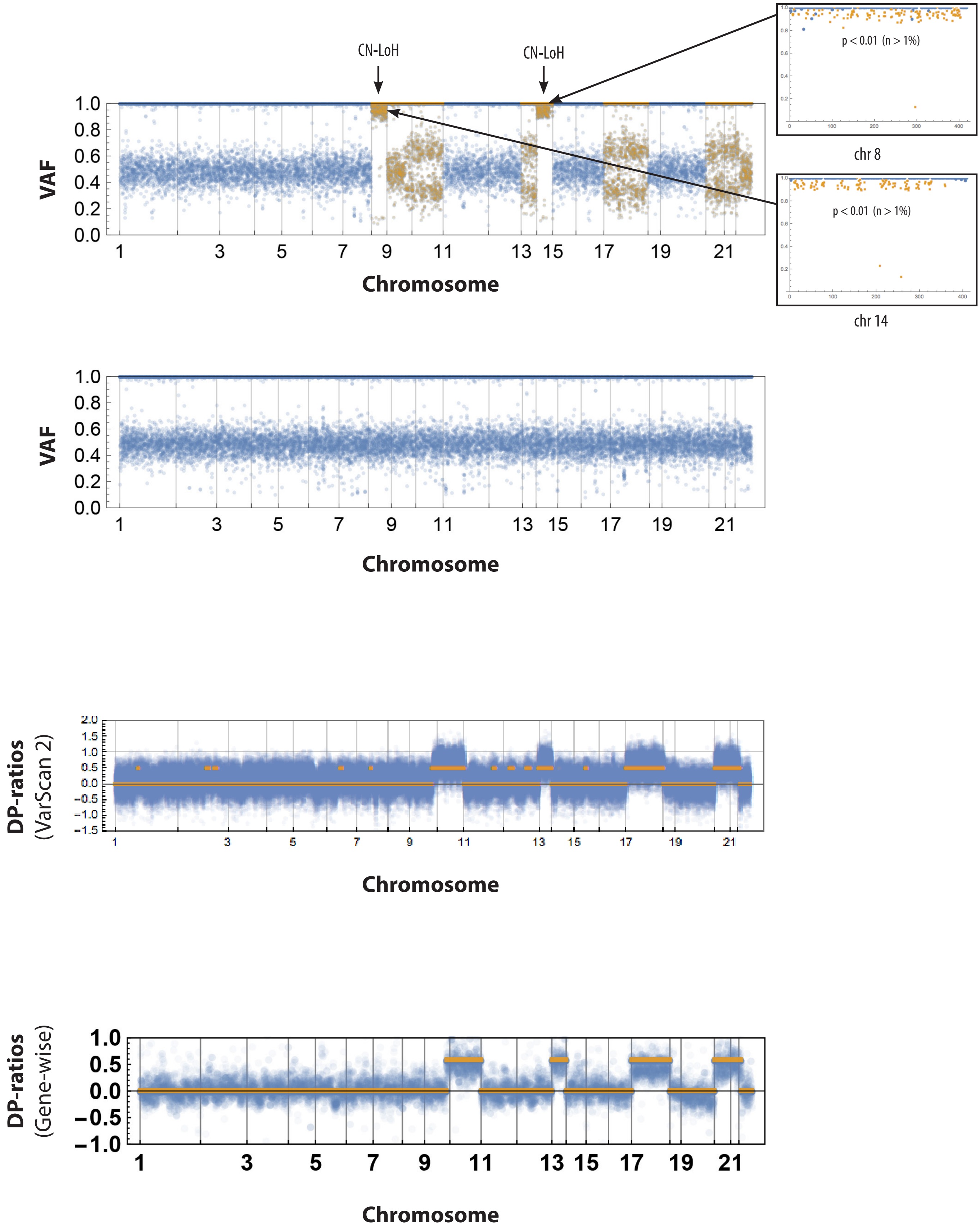

### Acute myeloid leukemia

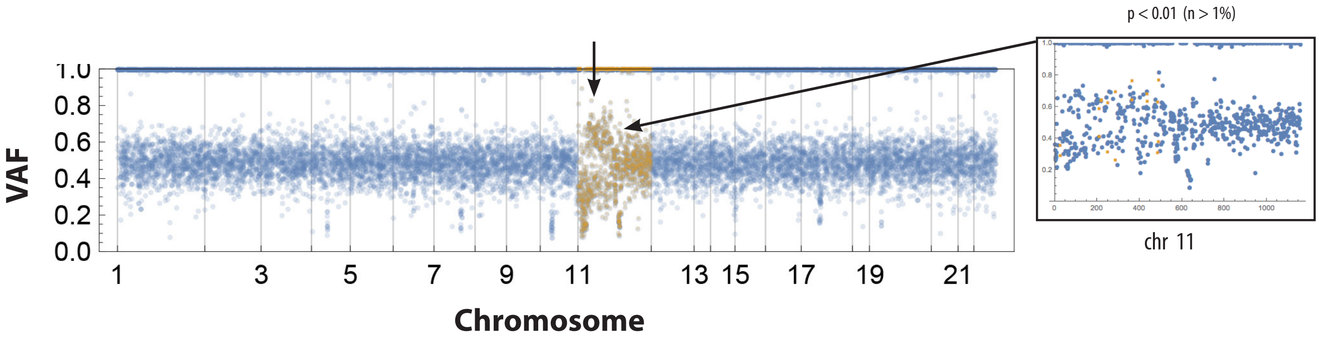


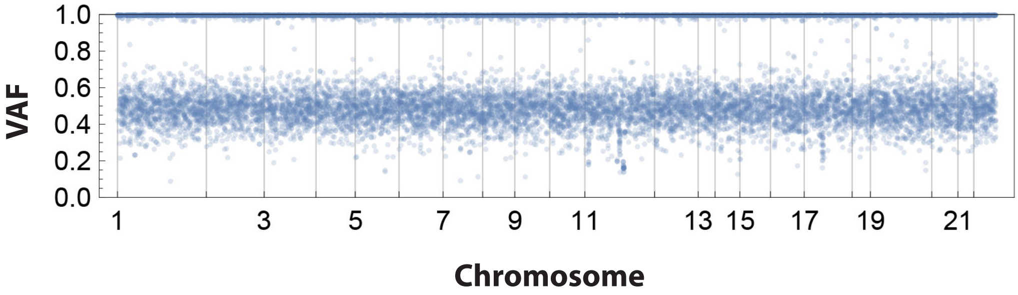


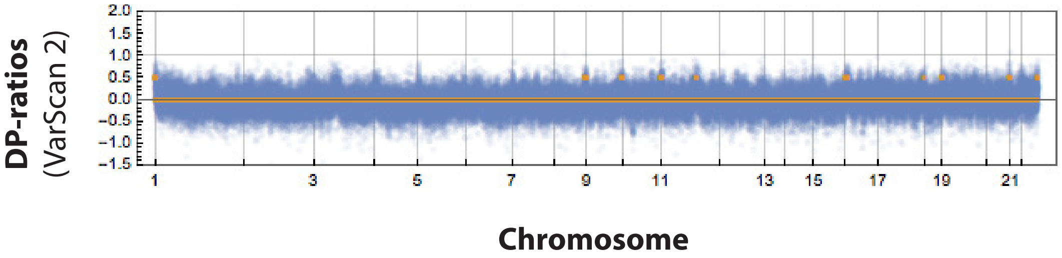


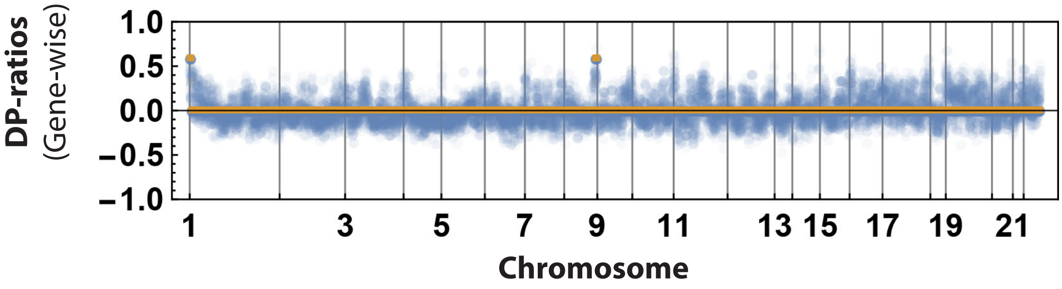
